# Structural investigation of QatB and QatC proteins in QatABCD anti-phage defense

**DOI:** 10.1101/2025.09.22.677948

**Authors:** Hyejin Oh, Euiyoung Bae

## Abstract

QatABCD is a widespread anti-phage defense system in prokaryotes comprising four protein components. QatC, a signature component, is homologous to QueC, an enzyme involved in nucleobase modification during queuosine biosynthesis. QatA and QatD are predicted to function as an ATPase and a nuclease, respectively, while QatB lacks identifiable sequence motifs. Here, we report the structural and functional characterization of QatB and QatC. We determined the structure of QatC bound to the ATP analog AMPPNP and performed structure-guided functional assays. We further found that QatB and QatC form a stable heterodimer and solved the structure of the QatB–QatC complex. In addition to unveiling the structure of QatB, structural analysis suggested that it may serve as a substrate of QatC, implicating a potential regulatory mechanism. These findings provide structural and functional insights into QatB and QatC, laying a foundation for understanding the molecular mechanism of the QatABCD system in phage defense.

## Introduction

Bacteriophages or phages are the most abundant biological entities on Earth (1). These viruses specifically infect bacteria, applying selective pressure on their hosts (2,3). The intense interplay between hosts and viruses has resulted in an evolutionary arms race, driving bacteria to evolve a wide array of sophisticated defense strategies (4). Several bacterial anti-phage defense mechanisms have been extensively studied, including restriction-modification and CRISPR-Cas systems (5,6). The protein components of these systems have been repurposed for biotechnological applications, such as restriction enzymes and Cas9-based genome editing tools (7,8).

Despite significant advances in our understanding of bacterial anti-phage defense mechanisms (9–22), the molecular basis of certain systems remains poorly characterized. In recent years, the development of systematic, high-throughput bioinformatic and experimental approaches has greatly accelerated the discovery of previously unrecognized bacterial anti-phage defense systems (23–26). Candidate systems have been identified through bioinformatic prediction, based on their genomic proximity to known defense genes or prophage-encoded diversity hotspots, followed by experimental validation of their anti-phage function (23,24,26). Alternatively, fosmid libraries derived from random genomic DNA fragments of diverse bacterial strains have been screened for anti-phage activity (25).

The QatABCD anti-phage defense system was first identified in *Escherichia coli* NCTC9009 (24). When introduced into another *E. coli* strain that naturally lacks this system, QatABCD conferred resistance to multiple bacteriophages, including P1, λ, and T3 (24). Furthermore, the QatABCD system from *E. coli* 46-1 provides protection against T4 phage infection (27), and the homologous system from *Pseudomonas aeruginosa* has been shown to resist phages of the *Pbunavirus* and *Casadabanvirus* genera (28). This defense system comprises four genes, namely *qatA*, *qatB*, *qatC* and *qatD* (24). The name ‘Qat’ is derived from the characteristics of its components and refers to a “QueC-like protein associated with ATPase and TatD DNase” (24).

QatA is predicted to function as an ATPase, as demonstrated by the loss of defense activity following a D210A mutation in the Walker B motif (24). Although QatB lacks recognizable conserved motifs, deletion of the *qatB* gene markedly impairs the system’s anti-phage function (24). QatC shows sequence similarity to QueC, an enzyme that catalyzes ATP-dependent conversion of 7-carboxy-7-deazaguanine (CDG) into 7-amido-7-deazaguanine and then to 7-cyano-7-deazaguanine in the queuosine biosynthetic pathway (24,29). QueC belongs to the PP-loop ATP pyrophosphatase family, which facilitates activation of a first substrate through AMPylation (also known as adenylylation), a process that releases pyrophosphate and enables a subsequent nucleophilic attack by a second substrate (30). QatC exhibits significantly reduced anti-phage activity when its predicted ATP-binding site is mutated (24). QatD shares similarity with TatD nuclease, and substitutions at its predicted catalytic residues results in reduced anti-phage activity of the QatABCD system (24). These results collectively indicate that all four components are essential for its defensive function against bacteriophage infection.

In this study, we structurally and functionally characterize QatB and QatC. The crystal structure of QatC reveals hallmark features of the PP-loop ATP pyrophosphatase family, with its conserved PP-loop motif binding the ATP analog AMPPNP. Based on this structure, we assessed the ATP pyrophosphatase and DNA-binding activities of QatC. We also found that QatB and QatC form a stable heterodimeric complex, supported by the crystal structure of the QatB–QatC complex. Notably, the N-terminal protruding loop of QatB inserts into the putative active site of QatC, suggesting that QatB may serve as its substrate, undergoing N-terminal modification. Together, our findings provide structural and functional insights into QatB and QatC and advance our understanding of how the QatABCD system contributes to prokaryotic defense against phage infection.

## Materials and Methods

### Cloning, expression and purification

The plasmid containing the four genes of the QatABCD system in *E. coli* NCTC9009 was obtained from Addgene, USA. Each gene was amplified by polymerase chain reaction and ligated into pET21a with a C-terminal (His)_6_ tag (Figure S1). *E. coli* BL21(DE3) cells harboring each construct were grown in lysogeny broth at 37°C until the optical density at 600 nm reached 0.7. Protein overexpression was induced by adding 0.5 mM isopropyl β-D-1-thiogalactopyranoside at 17°C for 16 h. The cells were harvested by centrifugation and resuspended in buffer [300 mM NaCl, 10% (w/v) glycerol, 5 mM β-mercaptoethanol, 30 mM imidazole, 20 mM tris(hydroxymethyl)aminomethane (Tris)-HCl pH 8.0]. After sonication and centrifugation, the supernatant was loaded onto a 5 mL HisTrap HP column (Cytiva, USA) pre-equilibrated with the same buffer. Following a wash step, bound proteins were eluted using a linear gradient of imidazole up to 450 mM. Further purification was performed by size-exclusion chromatography (SEC) using a HiLoad 16/600 Superdex 200 pg column (Cytiva), equilibrated with buffer [300 mM NaCl, 5% (w/v) glycerol, 2 mM 1,4-dithiothreitol (DTT), 20 mM 4-(2-hydroxyethyl)-1-piperazineethanesulfonic acid (HEPES) pH 8.0].

For co-expression of the QatB and QatC proteins, the *qatB* and *qatC* genes were cloned into pET21a with a C-terminal (His)_6_ tag and pET41a (no tag), respectively (Figure S1). *E. coli* BL21(DE3) cells were co-transformed with these constructs. Protein expression and purification of the resulting protein complex were carried out using the same procedure as for the individual proteins.

### Crystallization, data collection, and structure determination

The QatC protein was mixed with the ATP analog, adenylyl-imidodiphosphate (AMPPNP) at a molar ratio of 1:1.5, followed by incubation at 4°C for 1 h. Crystals of the AMPPNP-bound QatC were obtained at 20°C using the hanging-drop vapor diffusion method from a 296 µM solution of the QatC–AMPPNP complex in buffer [300 mM NaCl, 2 mM DTT, 5% (w/v) glycerol, 20 mM HEPES pH 8.0] mixed in equal volume with reservoir solution [16% (w/v) PEG20000, 0.1 M 2-(N-morpholino)ethanesulfonic acid pH 6.0].

To obtain the QatB–QatC complex, individually purified QatB and QatC proteins were mixed at a molar ratio of 1:1.2 and incubated at 4°C for 1 h. The mixture was loaded onto a HiLoad 16/600 Supderdex 200 pg column (Cytiva) equilibrated with buffer [300 mM NaCl, 5 % (w/v) glycerol, 2 mM DTT, 20 mM HEPES pH 8.0], and co-eluted fractions were pooled. Crystals of the QatB–QatC complex were obtained at 20°C using the sitting-drop vapor diffusion method from a 177 µM solution of the protein complex mixed in equal volume of reservoir solution [30% (w/v) SOKALAN CP42 (Molecular Dimensions, UK), 0.1 M Tris-HCl pH 8.5].

Crystals were cryoprotected in reservoir solution supplemented with 20% (w/v) glycerol and flash-frozen in liquid nitrogen. Diffraction data were collected at 100 K at the beamline 7A of the Pohang Accelerator Laboratory. Diffraction images were processed using XDS (31) and CCP4 (32). Structure models predicted using ColabFold v.1.5.5 (33) were employed for molecular replacement phasing in PHENIX (34). Final structures were completed through iterative rounds of manual model building in Coot (35) and refinement in PHENIX (34). The stereochemical quality of the final models was assessed using MolProbity (36).

### ATP pyrophosphatase activity assay

ATP pyrophosphatase activity was measured using the EnzCheck Pyrophosphatase Assay Kit (Thermo Fisher Scientific, USA), which detects free pyrophosphate in solution. Proteins (2 µM each) were pre-incubated at 25°C for 10 min in buffer (1 mM MgCl_2_, 1 mM DTT, 20 mM Tris-HCl pH 8.0) containing inorganic pyrophosphatase (0.03 unit), purine nucleoside phosphorylase (PNP; 1 unit), and 0.2 mM 2-amino-6-mercapto-7-methylpurine ribonucleoside (MESG). Reactions were initiated by adding 5 µL of 1 mM ATP and incubated at 25°C for 90 min. The absorbance at 360 nm was measured to quantify the amount of 2-amino-6-mercapto-7-methylpurine, which was produced by PNP from MESG in the presence of inorganic phosphate. Standard curves were generated using KH_2_PO_4_.

### Electrophoretic mobility shift assay (EMSA)

Proteins were incubated with DNA (0.5 µM) or RNA (2.0 µM) in 20 µL buffer (5 mM MgCl_2_, 1 mM DTT, 20 mM Tris-HCl pH 8.0) at 25°C for 30 min. Following the addition of 4 µL loading buffer [30 mM MgCl_2_, 6 mM DTT, 60% (w/v) glycerol, 0.1% (w/v) bromophenol blue, 120 mM Tris-HCl pH 8.0], the samples were subjected to native polyacrylamide gel electrophoresis (PAGE) and visualized by ethidium bromide staining.

### Analytical SEC

Analytical SEC was performed using a Superdex 200 Increase 10/300 GL column (Cytiva) pre-equilibrated with buffer [300 mM NaCl, 2 mM DTT, 5% (w/v) glycerol, 20 mM HEPES pH 8.0]. Samples (20 µM each) were incubated at 4°C for 1 h prior to injection and loaded onto the column at a flow rate of 0.5 mL/min. Elution fractions were analyzed by sodium dodecyl sulfate (SDS)-PAGE and visualized by Coomassie staining.

## Results

### Crystal structure of QatC reveals a conserved PP-loop core with variable surrounding regions

To gain structural insights into the anti-phage mechanism of the QatABCD system, we attempted to determine the crystal structures of individually purified Qat proteins. However, crystallization of QatA, QatB and QatC was initially unsuccessful. QatD protein was insoluble under our experimental conditions. Given that QatC has been predicted to share similarity with QueC, a member of the PP-loop ATP pyrophosphatase family (24), we attempted crystallization in the presence of the ATP analog AMPPNP. This approach has proven successful for several proteins in this family (30,37,38). Notably, we successfully determined the crystal structure of AMPPNP-bound QatC at a resolution of 3.1 Å (Figure 1A). Data collection, phasing, and refinement statistics are summarized in Table 1. The crystal structure revealed two QatC chains and two AMPPNP molecules in a single asymmetric unit. Structural alignment of the two chains yielded a root mean square deviation (RMSD) of 0.38 Å, suggesting that they share nearly identical conformations. Although two chains are present in the asymmetric unit, SEC analysis indicated that QatC exists as a monomer in solution (see below). Therefore, the dimer observed in the crystal structure is most likely a consequence of crystal packing rather than a biologically relevant oligomeric state. Accordingly, we hereafter describe chain A as the monomeric structure of QatC.

**Figure 1.**
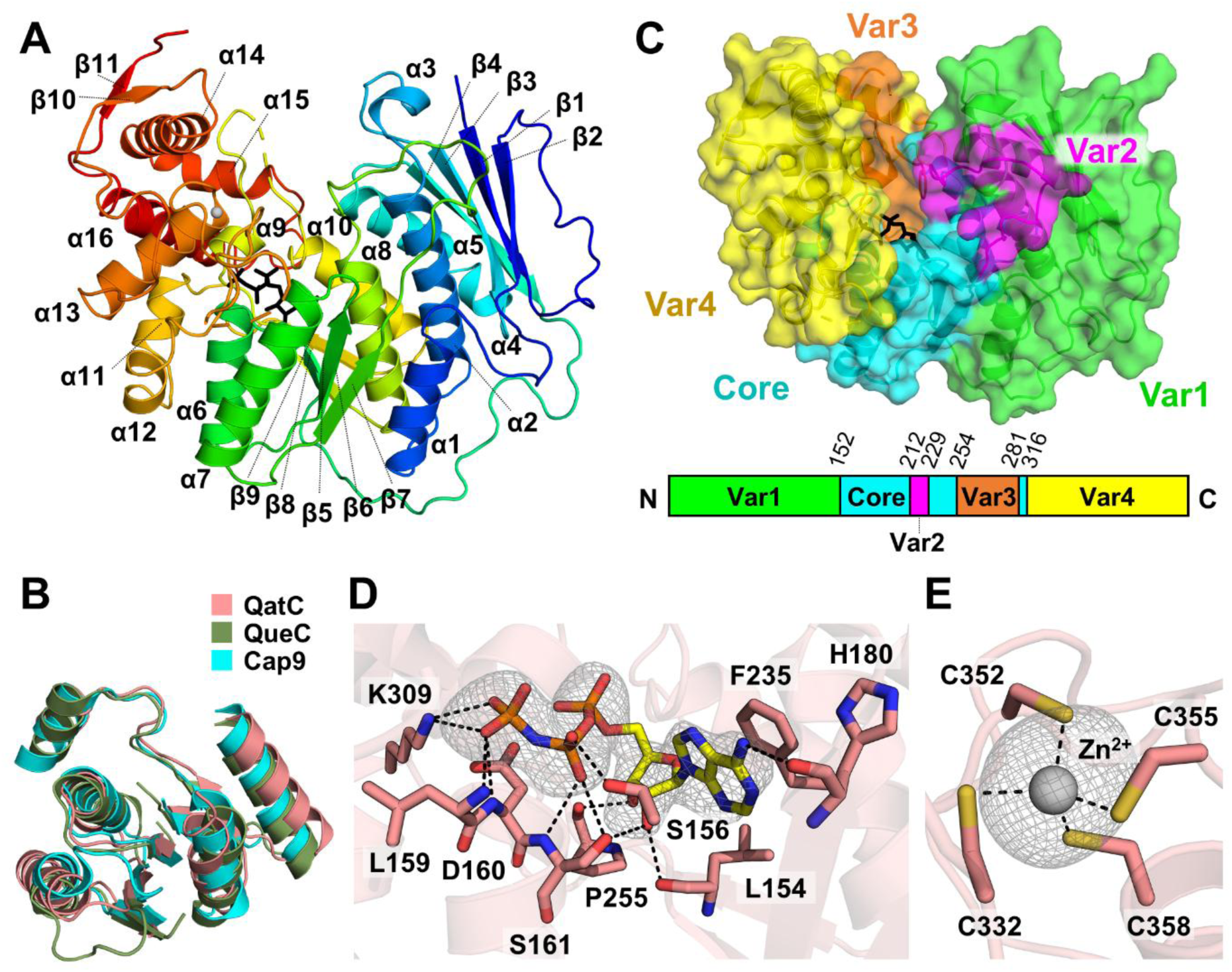
Crystal structure of QatC. (A) Overall structure of QatC. The protein is shown in a rainbow color scheme, from the N terminus (blue) to the C terminus (red). Secondary structural elements are annotated. The bound ATP analog, AMPPNP, is represented as black sticks. (B) Structural alignment of the conserved core domain of QatC (pink) with those of the PP-loop ATP pyrophosphatase family members QueC (olive) and Cap9 (cyan). (C) Conserved and variable regions in QatC. The structure exhibits a conserved Rossmann-like fold core domain with four variable regions (Var1–Var4). AMPPNP is depicted as black sticks. A schematic linear diagram of the domain organization is shown with residue numbers. (D) AMPPNP-binding site in QatC. The m*F*_obs_−D*F*_calc_ omit map is contoured at 3.0σ for AMPPMP. Surrounding QatC residues are also shown. (E) Zn-binding site in QatC. The Zn^2+^ ion is represented as a grey sphere. The m*F*_obs_−D*F*_calc_ omit map is contoured at 3.0σ for the Zn^2+^ ion. The coordinating Cys residues from QatC are also shown.

**Table 1.** Data collection, phasing and refinement statistics.

|  | QatC-AMPPNP<sup>a</sup> | QatB-QatC<sup>a</sup> |
| --- | --- | --- |
| Space group | P3<sub>1</sub> | I222 |
| Unit cell parameters (Å) | a=85.29 b=85.29 c=120.55<br>$\alpha=\beta=90^\circ$, $\gamma=120^\circ$ | a=115.41 b=119.59 c=120.17<br>$\alpha=\beta=\gamma=90^\circ$ |
| Solvent content (%) |  |  |
| Wavelength (Å) | 0.9793 | 0.9793 |
| **Data collection statistics** |  |  |
| Resolution range (Å) | 36.93-3.10 (3.31-3.10) | 30.00-2.58 (2.65-2.58) |
| Number of reflections | 187958 (33663) | 346896 (27813) |
| Completeness (%) | 100.0 (100.0) | 95.0 (99.9) |
| R<sub>merge</sub><sup>b</sup> | 0.362(0.989) | 0.417 (3.204) |
| CC1/2 | 0.965(0.814) | 0.995(0.712) |
| CC* | 0.991(0.947) | 0.999(0.912) |
| Redundancy | 10.5(10.4) | 13.8(13.1) |
| Mean I/$\sigma$ | 3.2(1.6) | 9.3(1.6) |
| **Refinement statistics** |  |  |
| Resolution range (Å) | 36.93-3.10 (3.31-3.10) | 30.00-2.58 (2.65-2.58) |
| R<sub>cryst</sub><sup>c</sup>/R<sub>free</sub><sup>d</sup> (%) | 17.0/25.2 | 19.7/27.8 |
| RMSD bonds (Å) | 0.011 | 0.009 |
| RMSD angles (deg) | 1.40 | 1.21 |
| Average B-factor (Å<sup>2</sup>) | 49.7 | 52.8 |
| Number of water molecules | 21 | 42 |
| Ramachandran favored (%) | 94.1 | 93.8 |
| Ramachandran allowed (%) | 5.6 | 5.9 |

<sup>a</sup>Values in parentheses are for the highest-resolution shell.
<sup>b</sup>R<sub>merge</sub> = $\sum_h \sum_i |I_i(h) - \langle I(h) \rangle| / \sum_h \sum_i I_i(h)$ , where $I_i(h)$ is the intensity of an individual measurement of the reflection and $\langle I(h) \rangle$ is the mean intensity of the reflection.
<sup>c</sup>R<sub>cryst</sub> = $\sum_h ||F_{obs}| - |F_{calc}|| / \sum_h |F_{obs}|$ , where $F_{obs}$ and $F_{calc}$ are the observed and calculated structure factor amplitudes, respectively.
<sup>d</sup>R<sub>free</sub> was calculated as R<sub>cryst</sub> using ~10% of the randomly selected unique reflections that were omitted from structure refinement.

The crystal structure of QatC reveals a Rossmann-like fold core (residues 152–211, 229–253, and 281–315), in which a five-stranded parallel β-sheet (β7-β6-β5-β8-β9) is sandwiched between two layers of α-helices: α6, α7, and α11 on one side, and α8 and α10 on the opposite side (Figure 1A, B). Similar to other members of the PP-loop ATP pyrophosphatase family, QatC possesses four additional variable regions (Var1–Var4) surrounding the conserved Rossmann-like core (Figure 1C). The N-terminal Var1 region (residues 1–151) comprises a four-stranded parallel β-sheet (β2-β1-β3-β4) with five α-helices (α1–α5) arranged along its concave side. The Var2 (residues 212–228) and Var3 (residues 254–280) regions, which primarily consist of flexible loops, are inserted within the Rossmann-like fold. The C-terminal Var4 region (residues 316–457) contains five α-helices (α12–α16).

Adjacent to the PP-loop motif (residues 156–161) in the Rossmann-like core of the QatC structure, we observed an additional electron density that was well accommodated with AMPPNP (Figure 1D). The binding mode and interacting residues closely resemble those observed in other PP-loop ATP pyrophosphatases. Phe235 engages in π-stacking with the adenine ring of ATP. The main chain oxygen of His180 forms a hydrogen bond with the N6 amino group of the adenine. The backbone oxygen of Leu154 and side chain hydroxyl group of Ser161 are hydrogen-bonded to the 2′-hydroxyl group of the ribose moiety, and the backbone oxygen of Pro255 engages with the 3′-hydroxyl group. Additionally, the side chain hydroxyl groups of Ser156 and Ser161, along with the backbone nitrogen of Ser161 interact with the β-phosphate of ATP, while the backbone nitrogen of Leu159 and Asp160 with the side chain of Lys309 participate in hydrogen bonding with the γ-phosphate. These structural observations indicate that the PP-loop motif and surrounding residues in QatC play a crucial role in ATP binding. Notably, a previous study demonstrated that the S156A/S161A mutations in QatC completely abolished the anti-phage protection conferred by the QatABCD defense system (24). Collectively, these findings suggest a direct link between ATP binding by QatC and the protective function of the anti-phage system.

Within the loop connecting α12 and α13 in the Var4 region, we identified a potential zinc-binding motif comprising four coordinating cysteine residues (Cys332, Cys352, Cys355, and Cys358) (Figure 1E). The electron density observed near this motif was consistent with a zinc ion, as modeling other ions or a water molecule did not fit the data well. This zinc binding motif is also conserved in QueC (Figure S2), where the zinc ion has been suggested to play a structural role (29).

### Structure-guided functional analysis of QatC

QatC was predicted to be a QueC-like protein (24), and the structural similarity search using the DALI server (39) revealed that QatC shares structural homology with members of the PP-loop ATP pyrophosphatase family (Table S1). PP-loop ATP pyrophosphatases catalyze a two-step reaction (Figure 2A). In the first step, ATP is hydrolyzed into AMP, and the adenylyl group is transferred to a substrate to generate an AMPylated intermediate (30). This is followed by a nucleophilic attack by a second substrate on the intermediate, which results in a covalent linkage between the two substrates and the release of AMP (30). A hallmark of this enzyme family is the conserved ATP-to-AMP hydrolysis mechanism. The presence of a conserved Rossmann-like PP-loop core domain in QatC, along with the observed binding of an ATP analog to this domain (Figure 1B, D), suggests that QatC may function as an ATP pyrophosphatase.

**Figure 2.**
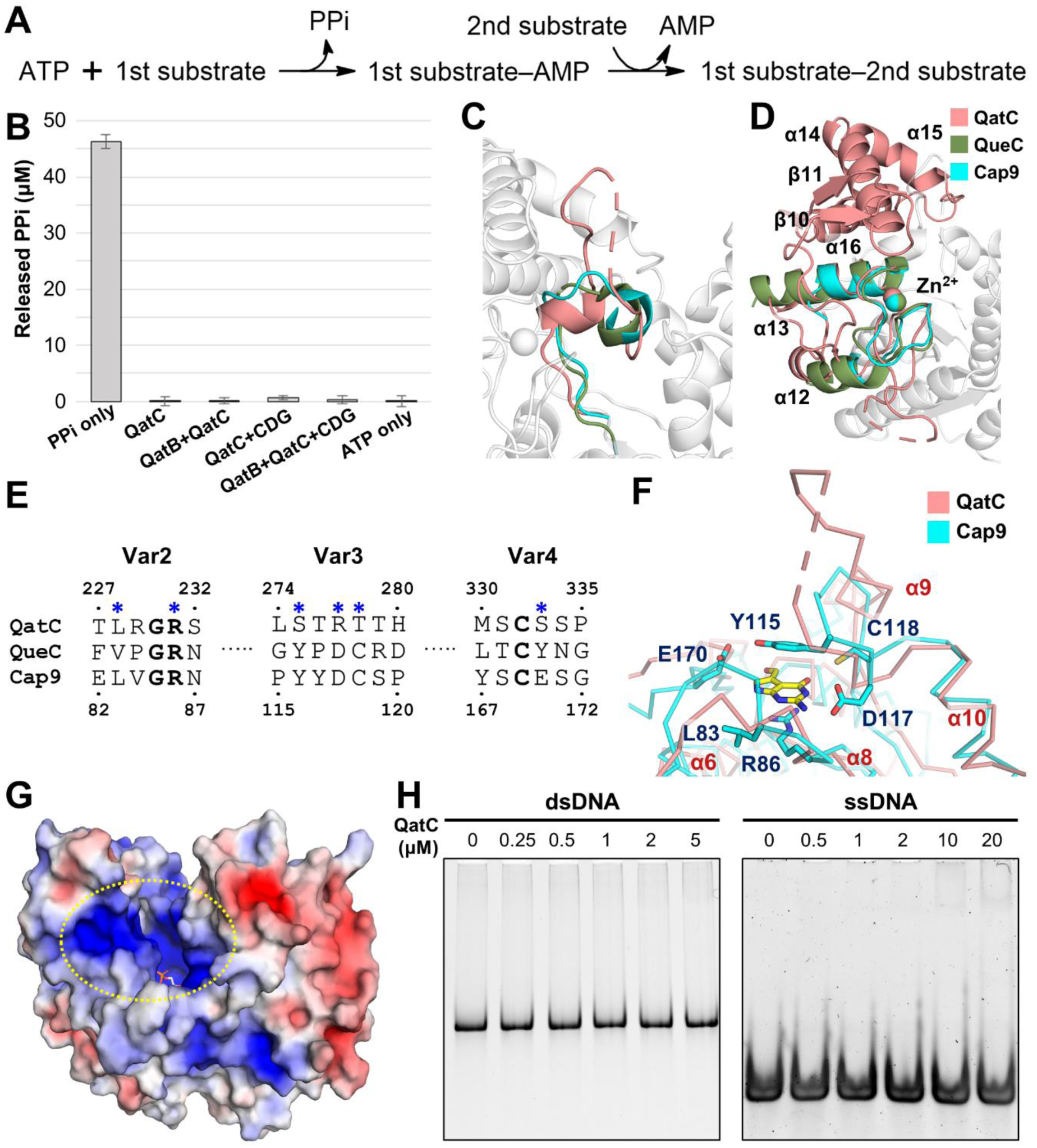
Functional assays of QatC. (A) Catalytic mechanism of PP-loop ATP pyrophosphatase family proteins. The first substrate is activated by AMPylation, followed by a nucleophilic attack by the second substrate. (B) ATP pyrophosphatase activity assay of QatC. The amount of released pyrophosphate from three independent experiments is shown as mean ± standard error of the mean. Inorganic pyrophosphate (PPi)-only and ATP-only samples were included as positive and negative controls, respectively. (C, D) Structural comparison of Var3 (C) and Var4 (D) regions of QatC (pink), QueC (olive), and Cap9 (cyan). QatC exhibits an extended loop in Var3 and additional C-terminal structural elements in Var4. The rest of the QatC structure are shown in white. (E) Sequence alignment of the substrate-binding regions of QatC, QueC and Cap9. Residue numbers at the top and bottom correspond to QatC and Cap9, respectively. Substrate-interacting residues in Cap9 are indicated by blue asterisks. (F) Structural comparison of the substrate binding site in QatC (pink) and Cap9 (cyan). The modified N-terminus of CdnD is represented by the yellow sticks. (G) Electrostatic potential surface of QatC. A positively charged patch adjacent to the ATP-binding site is indicated. PyMOL software (the PyMOL Molecular Graphics System, Version 2.0 Schrödinger, LLC.) was used with the Adaptive Poisson-Boltzmann Solver plugin to generate the surface (red = −5.0 kT,, blue = +5.0 kT). (H) DNA binding assay of QatC. Samples were analyzed by native PAGE. Uncropped gel images are provided in Figure S10.

To test whether QatC shares the functional characteristics of PP-loop ATP pyrophosphatases, we performed ATP pyrophosphatase assays (Figure 2B). When incubated with ATP alone, QatC showed negligible catalytic activity. This result suggests that ATP alone is insufficient to activate QatC and that the presence of an additional substrate, subject to AMPylation, may be required to stimulate ATP hydrolysis and catalysis. Notably, this requirement for a substrate to undergo AMPylation for ATP pyrophosphatase activity has been reported for other members of the PP-loop ATP pyrophosphatase family (38,40,41). The top two structural neighbors of QatC identified by the DALI server (39) were *Bacillus subtilis* QueC and *Rhizobiales* sp. Cap9 [cGAS/DncV-like nucleotidyltransferase (CD-NTase)-associated protein 9] (Table S1). Both QueC and Cap9 use CDG as a substrate for AMPylation (29,42). To investigate whether QatC exhibits similar substrate specificity, we tested its ATP pyrophosphatase activity in the presence of CDG. However, no significant catalytic activity was observed (Figure 2B), suggesting that CDG is not the cognate substrate for AMPylation by QatC.

To identify a potential substrate required for the ATP pyrophosphatase activity of QatC, we compared the structural features of QatC with those of other members of the PP-loop ATP pyrophosphatase family. Among the four variable regions (Var1–Var4) surrounding the conserved Rossmann-fold core, Var3 and Var4 are known to be involved in substrate binding in PP-loop ATP pyrophosphatases (30). Var3 is inserted within the Rossmann-fold core, and Var4 extends from its C-terminus. As these enzymes AMPylate a diverse range of substrates, both regions exhibit high structural variability (30). Structural comparison between QatC and other PP-loop ATP pyrophosphatases revealed no close matches in these regions. Notably, QatC exhibited substantial structural differences even from QueC and Cap9, its closest structural homologs identified by the DALI server (39) (Figure 2C, D). The Var3 region in QatC forms an extended loop of 27 residues, whereas the corresponding loops in QueC and Cap9 are considerably shorter, comprising only 13 and 12 residues, respectively. In the Var4 region, QatC shares only partial structural similarity with QueC and Cap9, including the Zn-coordinating site. Compared to both QueC and Cap9, QatC contains three additional α-helices in the C-terminal extension of this region. Moreover, key substrate-binding residues identified in QueC and Cap9, such as Asp131 in QueC and Asp117 in Cap9, are not conserved in QatC (Figures 2E, F and S2). These observations suggest that QatC may act on a previously unidentified type of substrate.

To further explore the potential function of QatC, we examined the characteristics of other components in the QatABCD system. Notably, QatD was previously predicted to be homologous to TatD nuclease (24), which has been implicated in DNA repair and other DNA-related processes (43,44). In several bacterial anti-phage systems, including restriction-modification and CRISPR-Cas systems, DNA nucleases are often found in association with other DNA-interacting proteins (10,13,14,45). This raises the possibility that the QatABCD system may operate through a DNA-based defense mechanism and that QatC could interact with DNA. Structural analysis of QatC revealed that the cleft leading to the ATP-binding pocket forms a positively charged surface (Figure 2G), which may facilitate interactions with negatively charged molecules. Similar positively charged surfaces are observed near the ATP-binding sites of other PP-loop ATP pyrophosphatases (Figure S3), where they are known to mediate binding to negatively charged substrates such as tRNA or various nucleotides (46–49). Based on these observations, we hypothesized that QatC might also interact with DNA. However, EMSAs revealed no detectable DNA-binding activity (Figure 2H), suggesting that QatC does not directly interact with DNA.

### QatC forms a stable heterodimeric complex with QatB

In many bacterial anti-phage systems, proteins interact to form multi-component complexes, and such interactions are often essential for their defensive activities against bacteriophages (10,12–14,22). Based on this, we investigated whether QatC forms a complex with other components of the QatABCD system (Figures 3 and S4). Because QatD was insoluble under our experimental conditions, we focused on testing interactions between QatC and the remaining components, QatA and QatB. In our analytical SEC experiments using individually purified protein components, QatB and QatC co-migrated as a single peak in the SEC column, eluting earlier than each protein alone, indicating the formation of a larger complex (Figure 3A). Furthermore, when His-tagged QatB was co-expressed with untagged QatC in *E. coli*, the two proteins co-eluted during purification, confirming the interaction (Figure 3B). These results collectively demonstrate that QatC forms a stable complex with QatB

**Figure 3.**
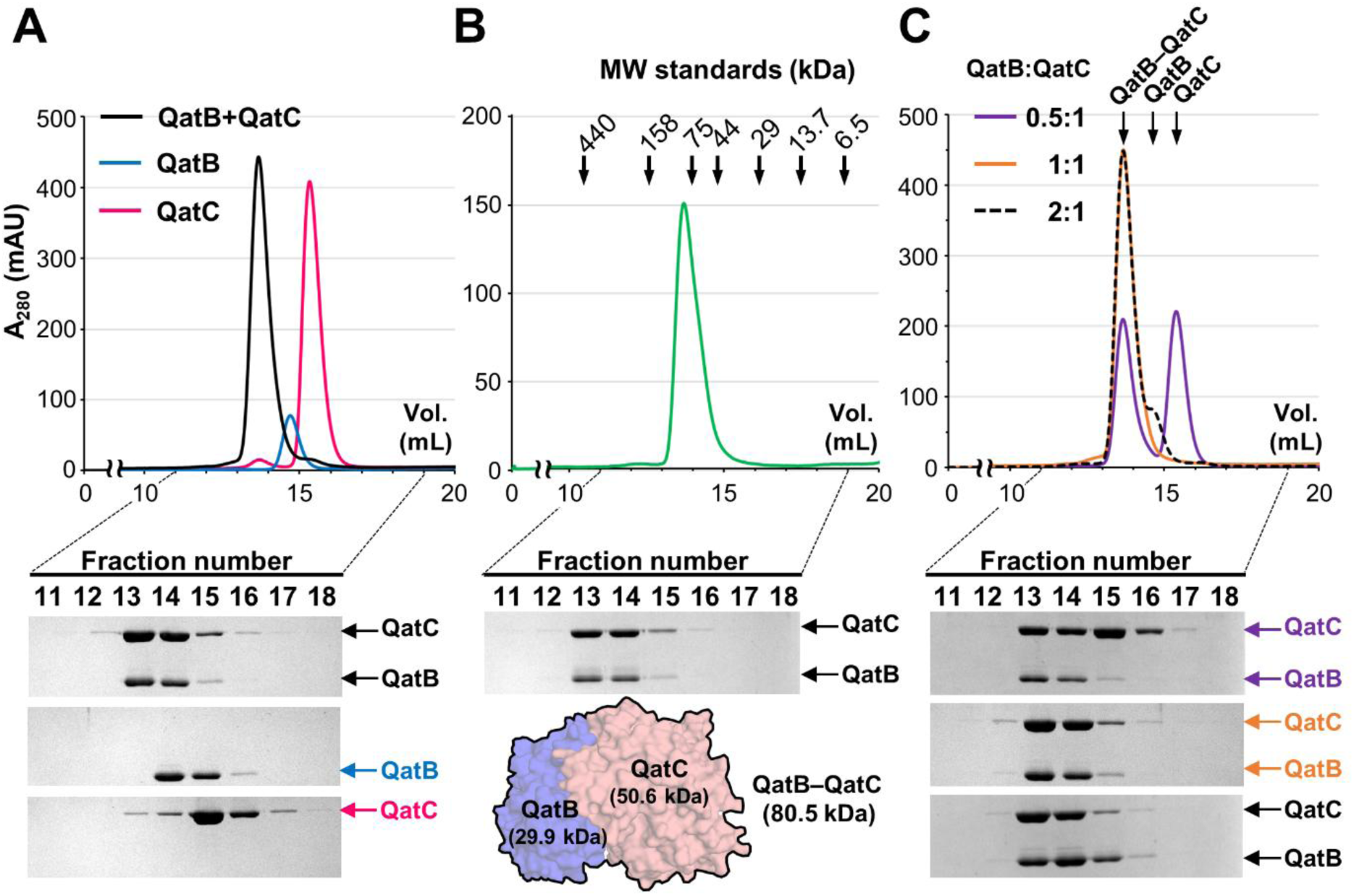
QatB and QatC form a heterodimeric complex. (A) Co-migration of individually purified QatB and QatC in SEC. QatB and QatC co-migrated (black) with a shorter retention time than that of QatB (blue) or QatC (pink) alone, indicating the formation of the QatB–QatC complex. (B) Co-elution of co-expressed QatB and QatC in SEC. When QatB and QatC were co-expressed in *E. coli*, they co-purified in SEC, further supporting complex formation. Theoretical molecular weights of QatB, QatC, and the QatB–QatC complex are indicated. (C) SEC analysis of the binding stoichiometry of the QatB–QatC complex. QatC was pre-incubated with increasing concentrations of QatB, and the samples were analyzed by SEC. The peak corresponding to the QatB–QatC complex increased until the QatB:QatC molar ratio reached 1:1 (purple and orange). Beyond this ratio, a second peak corresponding to excess QatB appeared, while the complex peak remained constant (black dashed line). Elution fractions were analyzed by SDS–PAGE. Uncropped gel images are provided in Figure S10.

To determine the stoichiometry of the QatB**–**QatC complex, we performed a series of analytical SEC experiments after pre-incubating a fixed amount of QatC with increasing concentrations of QatB (Figure 3C). As the amount of QatB increased, the peak corresponding to the complex gradually intensified. Once the molar ratio of QatB to QatC exceeded 1:1, a second peak corresponding to excess QatB appeared and increased in height, while the complex peak remained constant. These observations support a 1:1 binding ratio between QatB and QatC. According to the molecular weight estimated by SEC (Figure 3B), the complex likely exists as a heterodimer.

Given that QatC forms a stable complex with QatB, we then tested whether QatB affects the ATP pyrophosphatase activity or DNA-binding ability of QatC. However, neither activity was detected in the presence of QatB (Figures 2B and S5), suggesting that QatB is unlikely to serve as a substrate for AMPylation by QatC. Instead, the interaction with QatB may serve a different functional role in the QatABCD system.

### Crystal structure of QatB–QatC complex

To elucidate the molecular basis of the interaction between QatB and QatC, we determined the crystal structure of the QatB–QatC complex at a resolution of 2.58 Å (Figure 4A). Data collection, phasing, and refinement statistics are summarized in Table 1. The asymmetric unit contains one copy each of QatB and QatC. The complex was crystallized in the absence of AMPPNP. Nevertheless, a phosphate ion was modeled into the unaccounted electron density near the PP-loop motif of QatC, corresponding to the site of the terminal phosphate moiety observed in the AMPPNP-bound QatC structure (Figure S6). The phosphate ion is most likely of endogenous origin, derived from the *E. coli* expression system. Because QatC in the complex adopts a conformation essentially identical to the AMPPNP-bound structure (RMSD∼0.4 Å), the following structural description focuses on QatB and the interface between QatB and QatC.

**Figure 4.**
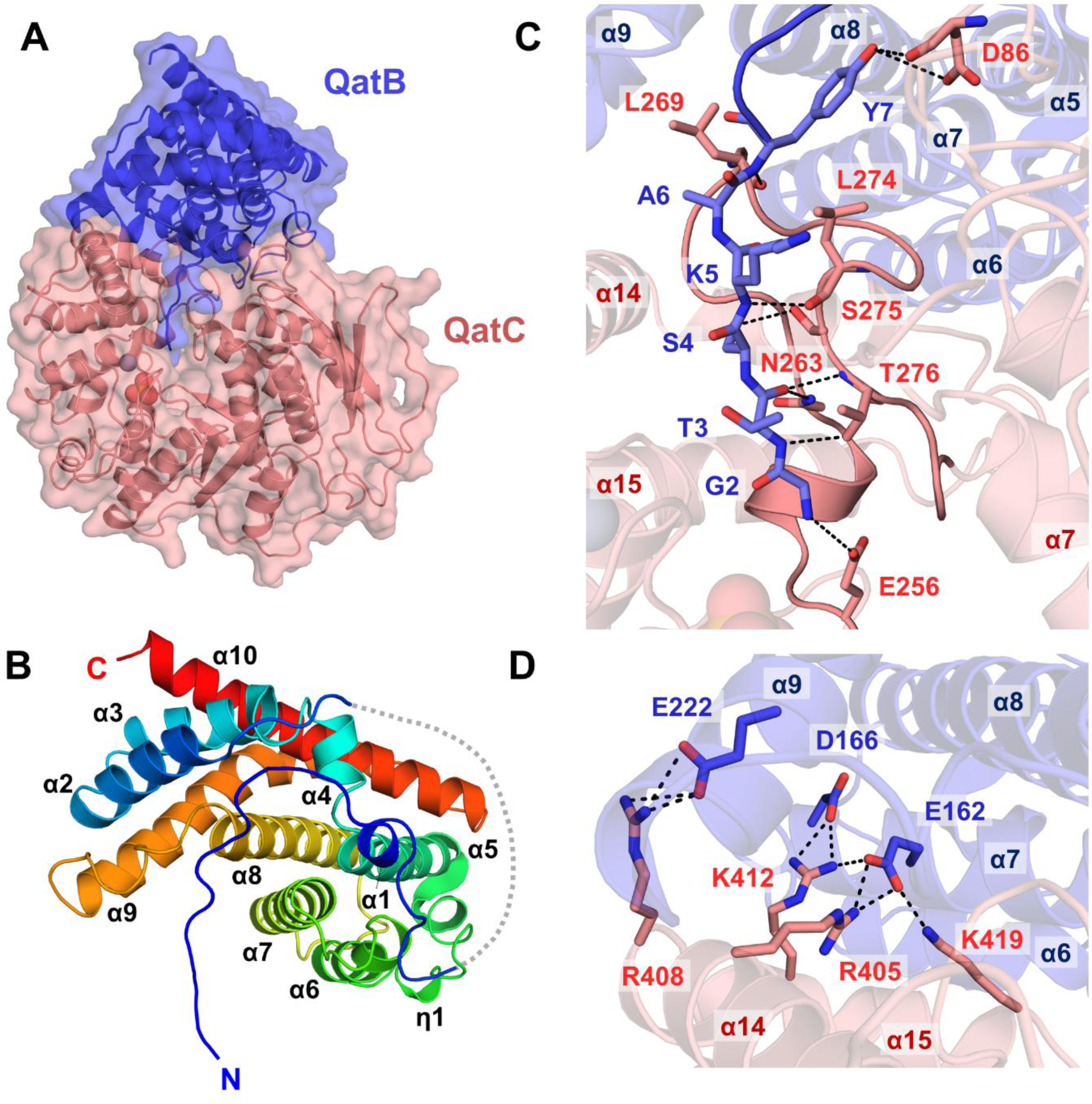
Crystal structure of the QatB–QatC complex. (A) Overall structure of the QatB– QatC complex. QatB (blue) and QatC (pink) form a heterodimeric assembly. (B) Structure of QatB. The structure is shown in a rainbow color scheme from the N terminus (blue) to the C terminus (red). Secondary structural elements are indicated. (C) Binding interface between the protruding N-terminal loop of QatB (blue) and QatC (pink). (D) Electrostatic interaction network between the main body of QatB (blue) and Var4 region of QatC (pink).

QatB exhibits an all-helical structure composed of 10 α-helices interspaced with loop regions (Figure 4B). Two long helices, α9 and α10, form a core of the structure in a bifurcated arrangement. On one side of this central pair are three α-helices (α2–α4), while a four-helix bundle (α5–α8) lies roughly perpendicular to the long helices. An intriguing feature of QatB is the presence of an extended N-terminal region including helix α1 and adjacent loop segments. This region wraps along the side of the four-helical bundle, with its N-terminal loop protruding outward from the main body of the protein. A structural similarity search using the DALI server (39) failed to identify any close homologs. Likewise, sequence-based BLAST searches (50) revealed no significant similarity beyond putative QatB proteins from other bacterial species.

The crystal structure of the QatB–QatC complex revealed a heterodimeric arrangement, consistent with the SEC results. Interface analysis using PISA (51) showed that the interaction buries 2360.6 Å^2^ and 2299.5 Å^2^ of solvent-accessible surface area on QatB and QatC, corresponding to 20.2% and 11.5% of their respective total surface areas. In addition, 13 salt bridges and 30 hydrogen bonds between QatB and QatC were identified. On the QatB side, the interface involves the protruding N-terminal loop (residues 2–10) and a region spanning from the C-terminus of α5 to the N-terminus of α7 (residues 136–225). On the QatC side, the interaction engages the conserved Rossmann-like core and three variable regions (Var1, Var3, and Var4). The N-terminal loop of QatB inserts into an extended cleft formed primarily by Var3 and Var4 of QatC, where it interacts mainly through van der Waals contacts and backbone-mediated hydrogen bonds (Figure 4C). A network of electrostatic interactions is also observed between the main body of QatB and Var4 of QatC (Figure 4D). In this interface, negatively charged residues in QatB (Glu162, Asp166, and Glu222) form salt bridges with positively charged residues in QatC (Arg405, Arg408, Arg412, and Lys419).

The most notable feature of the QatB–QatC complex structure is the insertion of the extended N-terminal segment of QatB into the putative active site in QatC (Figure 5A), suggesting that QatB may participate in the reaction catalyzed by QatC. This structural arrangement is reminiscent of the crystal structure of the CdnD–Cap9 complex (Figure 5B), in which Cap9, one of the closest structural homologs of QatC, forms a 2:2 complex with CdnD, a CD-NTase enzyme in the *Rhizobiales* type IV cyclic oligonucleotide-based antiviral signaling system (CBASS) (42). In the CdnD–Cap9 complex structure, the N-terminal loop of CdnD, which lacks the initiating Met1 residue due to post-translational processing, is inserted into the active site of Cap9 (42). Notably, the Gly2 residue of CdnD is conjugated to a 7-deazaguanine nucleobase (42). Based on this and other supporting evidence, it has been proposed that Cap9 catalyzes the modification of the CdnD N-terminus through a QueC-like mechanism, wherein CDG is first activated by AMPylation, followed by nucleophilic attack by the N-terminal amine of CdnD to complete the protein-nucleobase conjugation (42).

**Figure 5.**
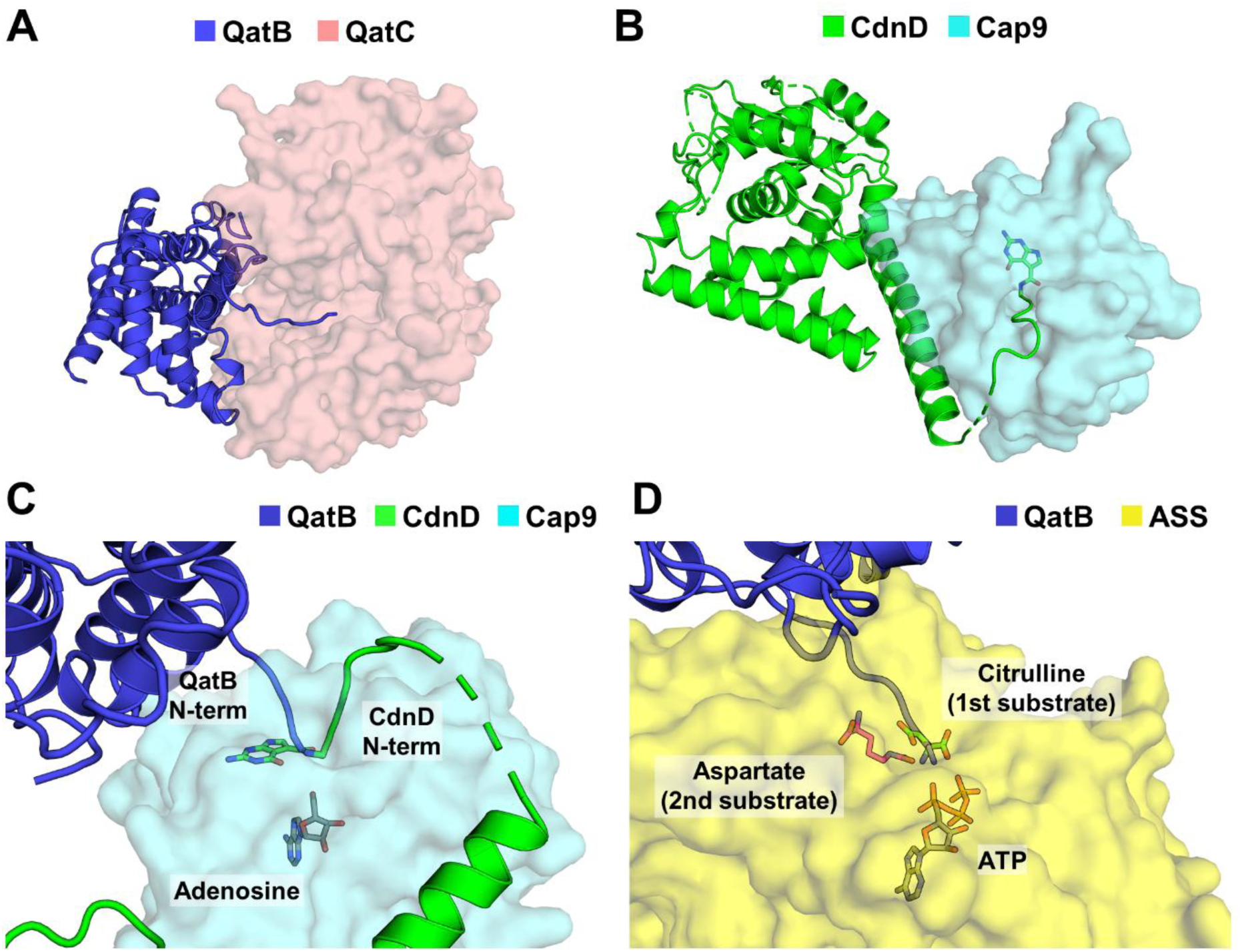
QatB functions as the putative second substrate of QatC. (A, B) Structural comparison of the QatB–QatC complex (A) and the CdnD–Cap9 complex (B; PDB ID: 9NTO). In both complexes, the extended N-terminal segment of one protein (QatB or CdnD) inserts into the active site of the other (QatC or Cap9). QatB, QatC, CdnD and Cap9 proteins are shown in blue, pink, green, and cyan, respectively. (C, D) Structural alignment of the putative active site of QatC with those of Cap9 (C) and argininosuccinate synthase (ASS) (D; PDB ID: 1K97). Upon alignment of the QatB–QatC complex with the CdnD–Cap9 complex and substrate-bound ASS, the N-terminal amine of QatB spatially overlaps with that of CdnD (C) and the second substrate of ASS (D). For clarity, QatC is omitted and only Cap9 and ASS are shown.

Despite the lack of overall structural similarity between QatB and CdnD, both exhibit a shared mode of active-site engagement with their QueC-like partners via long N-terminal tails (Figure 5A, B). Notably, removal of the initial Met1 is a post-translational modification in bacterial proteins, typically occurring when the second residue is a small amino acid such as glycine (52). Gly2 is conserved across both QatB and CdnD homologs (42) (Figure S7). In our QatB–QatC structure, Met1 was not modeled due to insufficient electron density. Structural alignment reveals that the N-terminal amines of Gly2 in QatB and CdnD are positioned in close proximity (∼1.2 Å apart) (Figure 5C). Moreover, superposition with argininosuccinate synthase (ASS), another member of the PP-loop ATP pyrophosphatase family, whose structure was determined in complex with ATP and its two substrates (38), reveals the N-terminus of Gly2 in QatB spatially overlaps with the position of the second substrate in ASS (Figure 5D). These findings support the notion that QatB serves as the second substrate of QatC, which first AMPylates an unidentified substrate that is subsequently attacked by the N-terminus of QatB to form a covalent linkage. The identity of the first substrate for QatC-mediated AMPylation, however, remains to be determined.

## Discussion

QatC exhibits structural similarity to QueC, including a conserved Rossmann-like fold core and a zinc binding motif, thereby supporting its previous annotation as a QueC-like protein (24). The central Rossmann-like domain is a hallmark of the PP-loop ATP pyrophosphatase family, to which both QatC and QueC belong (30). Members of this protein family typically employ a conserved catalytic mechanism that involves AMPylation of a first substrate, followed by a nucleophilic attack by a second substrate on the AMPylated intermediate, resulting in covalent linkage of the two components (30). Despite this shared mechanism, the specific reactions catalyzed by different family members are highly diverse (30). Notably, the identity of the AMPylated substrate varies widely, encompassing a broad range of molecules such as amino acids, nucleobases, and nucleotides (29,53–56). The diversity of first substrates in the PP-loop ATP pyrophosphatase family is primarily attributed to structural variations in the two variable regions (Var3 and Var4), which differ among individual family members (30).

Structural comparison of QatC with QueC and Cap9 revealed that the secondary structures and substrate-binding residues within the Var3 and Var4 regions are not conserved (Figure 2). Consistently, QatC did not exhibit ATP pyrophosphatase activity in the presence of CDG, the first substrate of both QueC and Cap9 (29,42). Furthermore, comparative analysis with other PP-loop ATP pyrophosphate family members with known structures and substrates did not yield sufficient clues to identify the first substrate of QatC, suggesting that it may use a previously uncharacterized molecule. One possibility is that the first substrate recognized by QatC is linked to a phage infection sensing mechanism. Given the remarkable diversity of first substrates within the PP-loop ATP pyrophosphatase family (30), the potential substrate undergoing AMPylation by QatC may include phage-derived factors, signaling molecules or their precursors, or intracellular metabolites that reflect the physiological state of the host cell (45,57–59).

Based on our structural data and other supporting evidence, we propose that QatB serves as the second substrate of QatC, participating in the nucleophilic attack on an unidentified first substrate. In the QatB–QatC complex, the protruding N-terminal loop of QatB inserts into the catalytic pocket of QatC. A similar binding mode involving a QueC-like protein is observed in the type IV CBASS system of *Rhizobiales* sp., where the N-terminal amine of CdnD penetrates the active site of Cap9 and is covalently conjugated to a 7-deazaguanine nucleobase derived from CDG, the first substrate of Cap9 (42). CBASS are widespread and diverse antiviral defense mechanisms in prokaryotes (60). They rely on cyclic oligonucleotides synthesized by CD-NTases, to activate effector proteins that trigger cell suicide (60). They are categorized into four types, each with distinct ancillary Cap proteins that regulate defense activities (60). In the type IV CBASS of *Rhizobiales* sp., Cap9 and Cap10 act as essential components along with the CD-NTase enzyme CdnD and its effector protein (42). Cap10 is homologous to tRNA-guanin transglycosylase, an enzyme involved in queuosine biosynthesis (61). A previous study proposed that Cap9 mediates nucleobase conjugation at the N-terminus of CdnD, while Cap10 binds the modified CdnD and facilitates its transfer to an unidentified target (42).

The QatABCD system and the type IV CBASS share several common features, including the presence of a QueC-like component and its unique binding mode with the long N-terminal loop of a partner protein. Therefore, it is tempting to hypothesize that the QatABCD may employ an anti-phage mechanism similar to that of CBASS. However, there are substantial differences between the two systems that preclude a straightforward mechanistic comparison. Despite the conserved structural features in the two QueC-like proteins, their binding partners, QatB and CdnD, do not exhibit any structural similarity beyond the presence of a protruding N-terminal loop (Figure S8), making it unlikely that QatB functions in cyclic oligonucleotide production as CdnD does. Further comparison of other components between the two systems also revealed clear differences. They share no significant similarity at either the sequence or structural level. Interestingly, QatD, which is related to TatD nuclease, is predicted to adopt a TIM barrel fold as observed in Cap10 (42) (Figure S9). However, this structural fold is associated with a wide range of enzymatic activities including isomerase, aldolase, and triesterase functions, making it insufficient to support any specific functional inference (43,62,63). Notably, all of the catalytic residues required for TatD nuclease activity are conserved in QatD, and mutating these residues markedly reduces the defensive function of the QatABCD system (24). This strongly suggests that QatD functions as a nuclease rather than as a Cap10-like transferase.

Given that the Cap proteins generally function as regulators of CD-NTase activity (60), the interaction between QatB and QatC may similarly play a role in modulating the defense function of the QatABCD system. QatC-mediated modification at the N-terminus of QatB may influence its catalytic activity and/or binding affinity to other components or unidentified targets. In our SEC analysis, QatB did not show detectable interaction with QatA (Figure S4). Nevertheless, we cannot rule out the possibility that the modified QatB, through its remodeled N-terminus, associates with other proteins similar to the interaction between the modified CdnD and Cap10 observed in the type IV CBASS (42). Due to the poor solubility of QatD under our experimental conditions, we were unable to assess its interaction with other components. Further structural and functional investigations are required to elucidate the precise roles of each protein in the QatABCD system and to define its overall mechanism of anti-phage defense.

## Data availability

The atomic coordinates and structure factors were deposited in the Protein Data Bank under the accession numbers 9VLK and 9VLL for AMPPNP-bound QatC and the QatB–QatC complex, respectively.

## Acknowledgments

We thank the staff of beamline 7A of the Pohang Accelerator Laboratory for their support with data collection.

## Author contributions

E.B. conceived the study. H.O. performed the protein purification, crystallization, functional assays, and SEC. H.O. and E.B. determined the crystal structures, analyzed the data, and wrote the manuscript.

