## Supplementary Data for "Structural investigation of QatB and QatC proteins in QatABCD anti-phage defense"

**Table S1. Top 10 structural neighbors of QatC identified by the Dali server (39).**

| Protein | Organism | PDB ID | Z-score | RMSD (Å) | Number of superposed residues |
|---|---|---|---|---|---|
| QueC | *Bacillus subtilis* | 3BL5 | 14.4 | 2.9 | 177 |
| Cap9 | *Rhizobiales* sp. | 9NTO | 13.8 | 2.4 | 168 |
| QueC | *Erwinia carotovora* subsp. atroseptica SCRI1043 | 2PG3 | 13.5 | 3.2 | 186 |
| ThiI | *Thermotoga maritima* MSB8 | 4KR6 | 10.1 | 3.2 | 156 |
| Argininosuccinate synthase | *Thermotoga maritima* | 1VL2 | 9.8 | 3.7 | 155 |
| LarE | *Methanococcus maripaludis* | 8CNZ | 9.2 | 3.2 | 157 |
| ThiI | *Bacillus anthracis* str. Ames | 2C5S | 9.2 | 3.3 | 147 |
| Hypothetical protein PH1313 | *Pyrococcus horikoshii* OT3 | 1VBK | 9.0 | 3.1 | 119 |
| NcsA | *Methanococcus maripaludis* S2 | 6SCY | 8.8 | 3.7 | 169 |
| TtuB | *Thermus thermophilus* HB27 | 5ZTB | 8.7 | 4.2 | 159 |

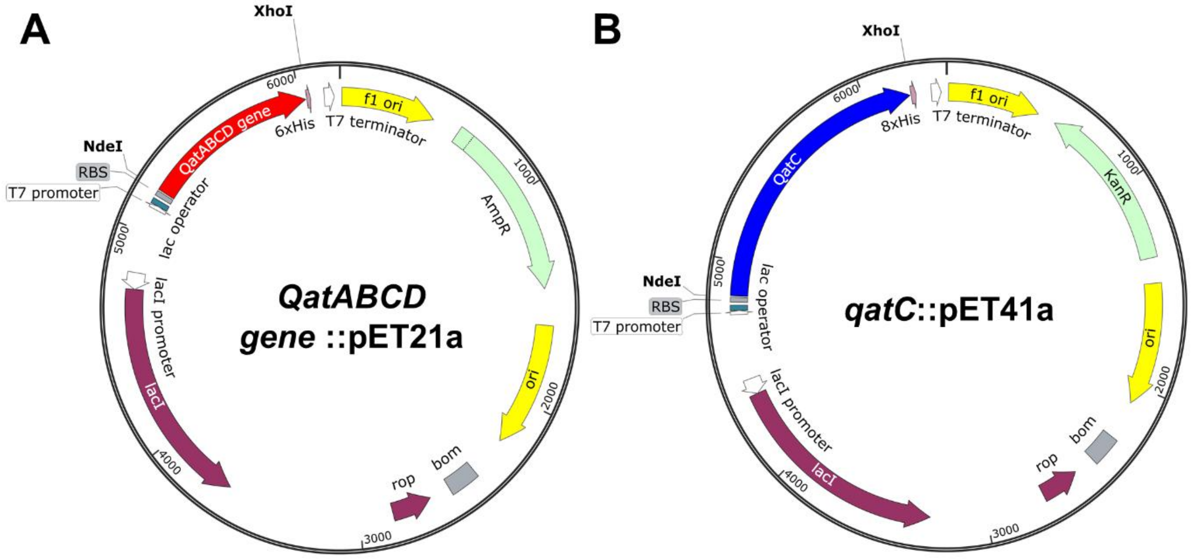


**Figure S1. Plasmid maps of expression vectors for QatABCD proteins.** Maps were generated by using SnapGene software (v.5.3.1).

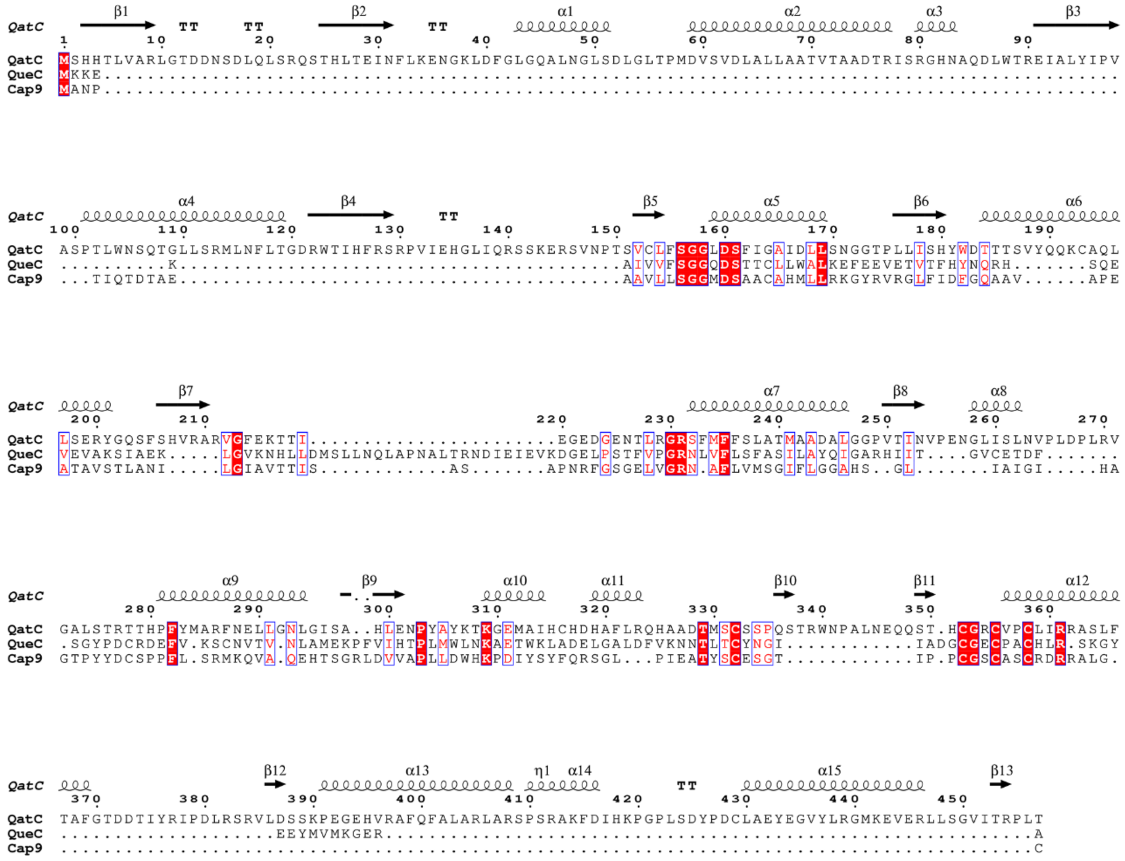


**Figure S2. Sequence alignment of QatC with QueC and Cap9.** White characters in red boxes indicate strict identity. Red characters represent similarity. The secondary structural elements in QatC are shown.

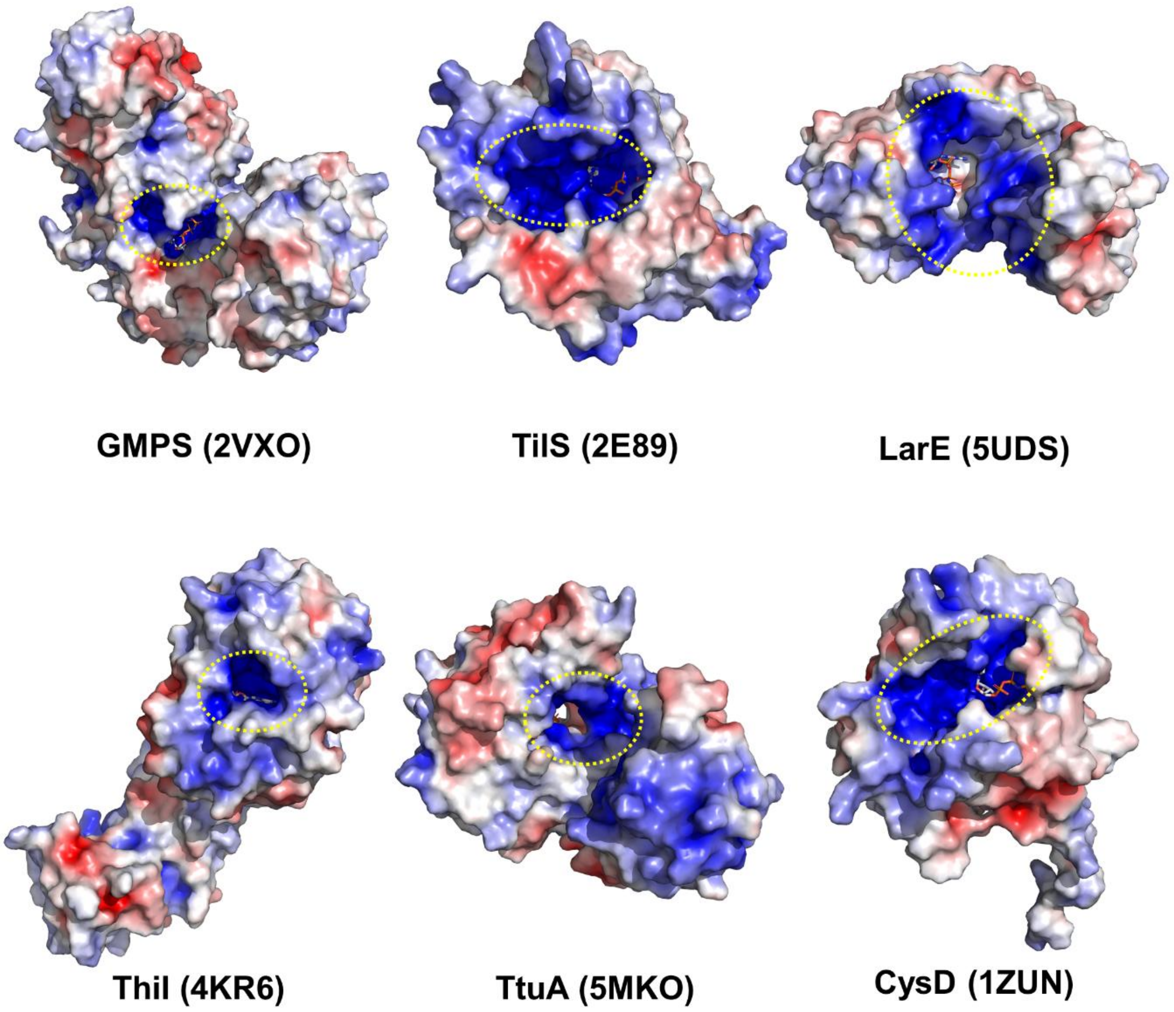


**Figure S3. Electrostatic potential surfaces of PP-loop ATP pyrophosphatase family proteins.** A positively charged patch adjacent to the ATP-binding site is indicated. PyMOL software (the PyMOL Molecular Graphics System, Version 2.0 Schrödinger, LLC.) was used with the Adaptive Poisson-Boltzmann Solver plugin to generate the surface (red = −5.0 kT, blue = +5.0 kT). The proteins shown include GMP synthetase (GMPS), tRNA$^{Ile}$-lysidine synthetase (TilS), a sulfur transferase involved in cofactor biosynthesis for lactate racemase (LarE), tRNA 4-thiouridine synthetase (ThiI), tRNA 5-methyl-2-thiouridine synthetase (TtuA), and sulfate adenylyltransferase subunit 2 (CysD).

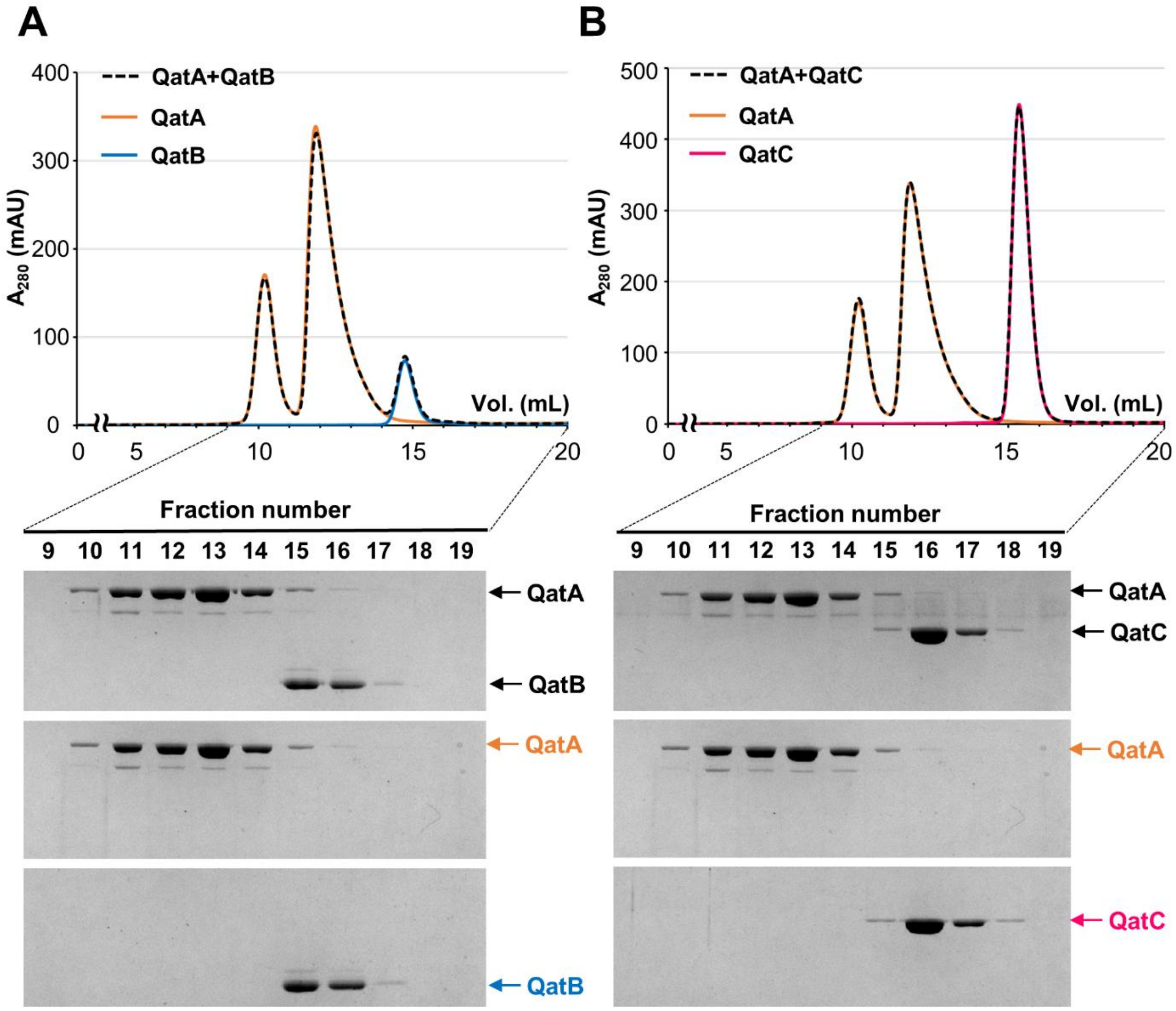


**Figure S4. SEC analysis for testing of the interactions between protein components of the QatABCD system.** QatA was pre-incubated with either QatB (A) or QatC (B), and the mixtures were analyzed by SEC. The chromatograms of the mixtures (black dashed lines) closely match the sum of the individual protein profiles, indicating no detectable interaction. Uncropped gel images are provided in Figure S10.

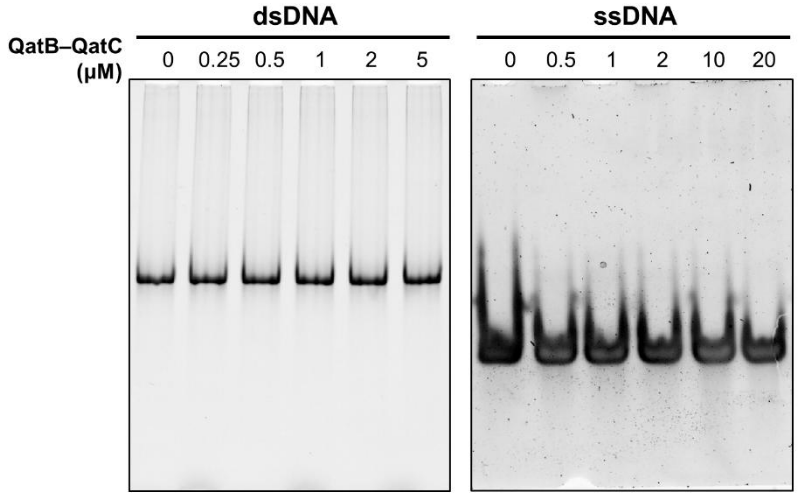


**Figure S5. DNA binding assay of the QatB–QatC complex.** Samples were analyzed by native PAGE. Uncropped gel images are provided in Figure S10.

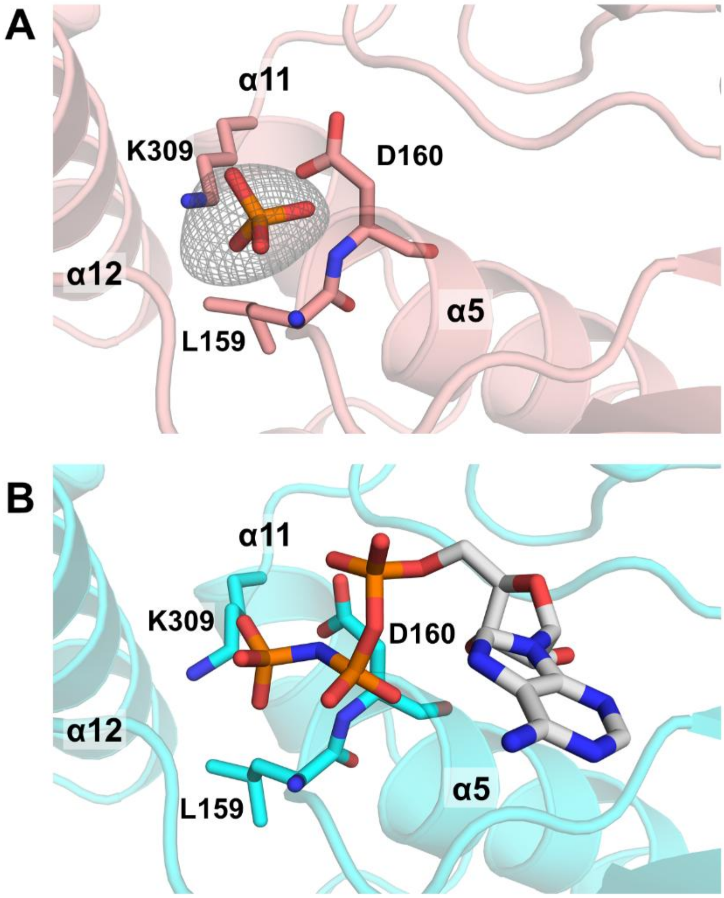


**Figure S6. A phosphate ion is observed near the PP-loop motif of QatC in the QatB–QatC complex structure.** (A) The phosphate ion and surrounding QatC residues in the structure of the QatB–QatC complex. The m$F_{obs}$–D$F_{calc}$ omit map is contoured at 3.0σ for the phosphate ion. (B) AMPPNP and surrounding QatC residues in the AMPPNP-bound QatC structure. The position of the phosphate ion in the QatB–QatC complex closely matches that of the terminal phosphate of AMPPNP.

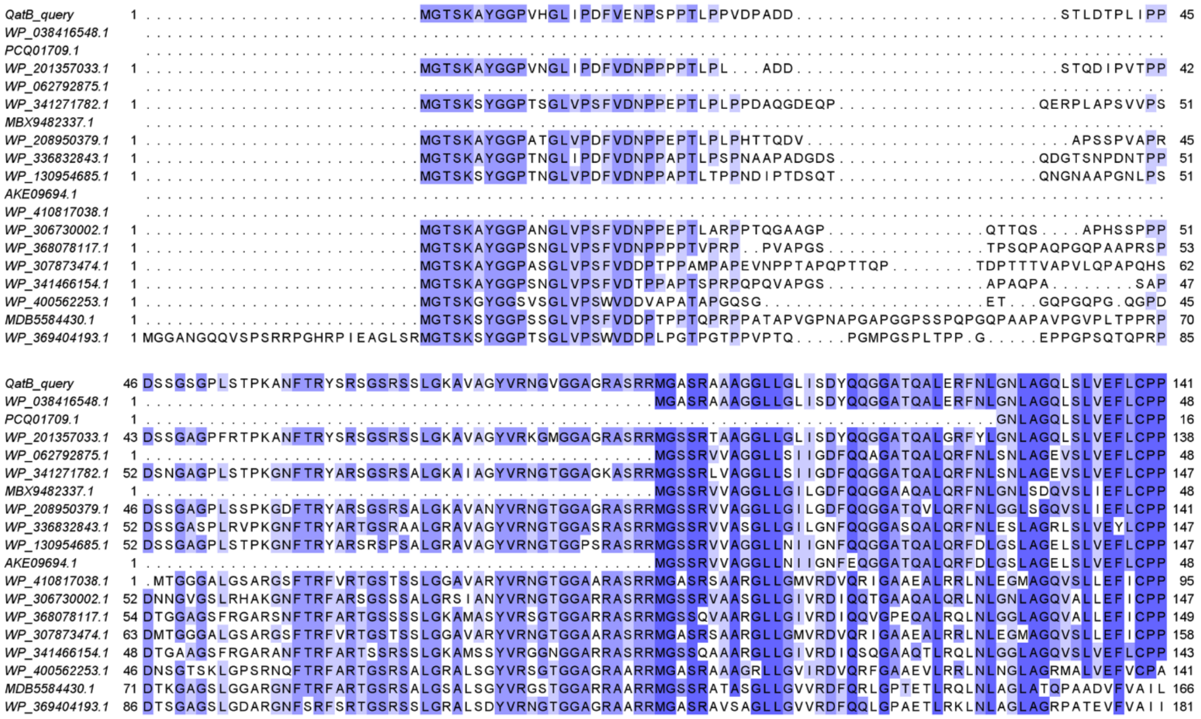

**Figure S7. Sequence alignment of the N-terminal regions of QatB homologs.** Amino acid conservation is indicated by blue shading.

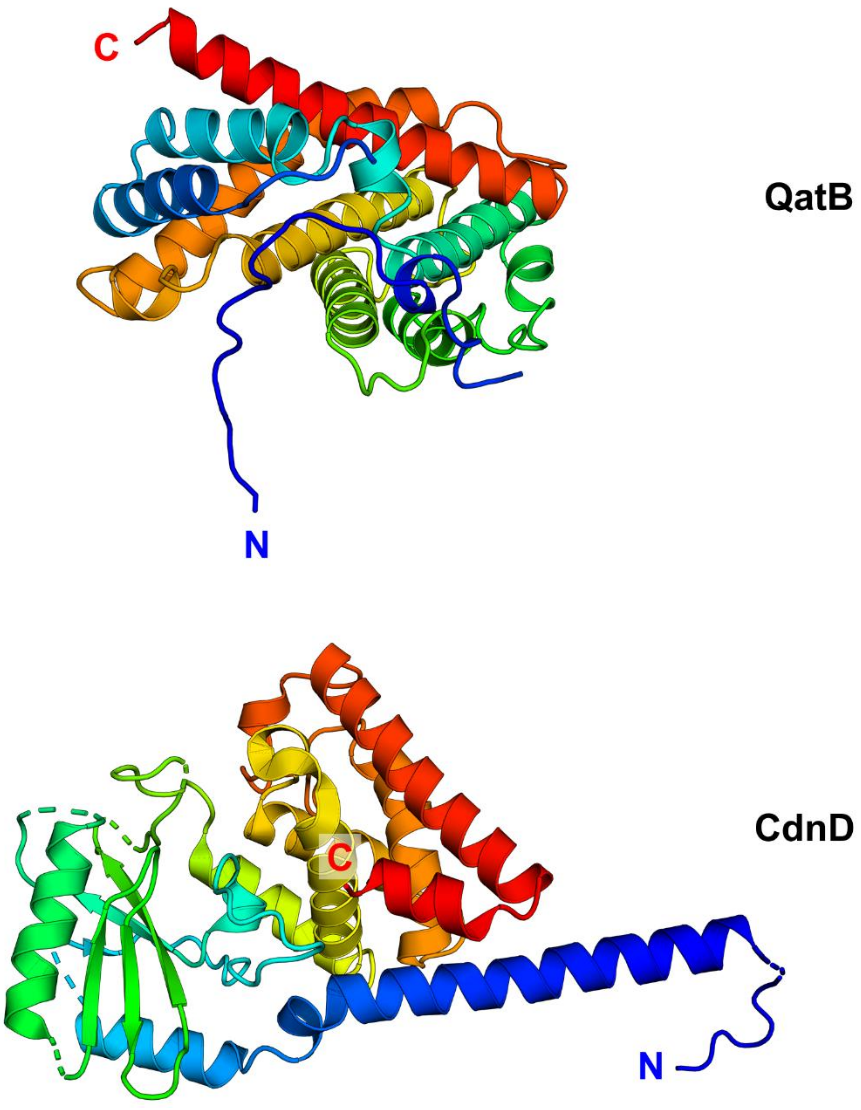


**Figure S8. Structural comparison of QatB and CdnD.** QatB (top) and CdnD (bottom) exhibit distinct overall structures, with the exception of the N-terminal protruding loop. The structures are shown in a rainbow format from the N terminus (blue) to the C terminus (red).

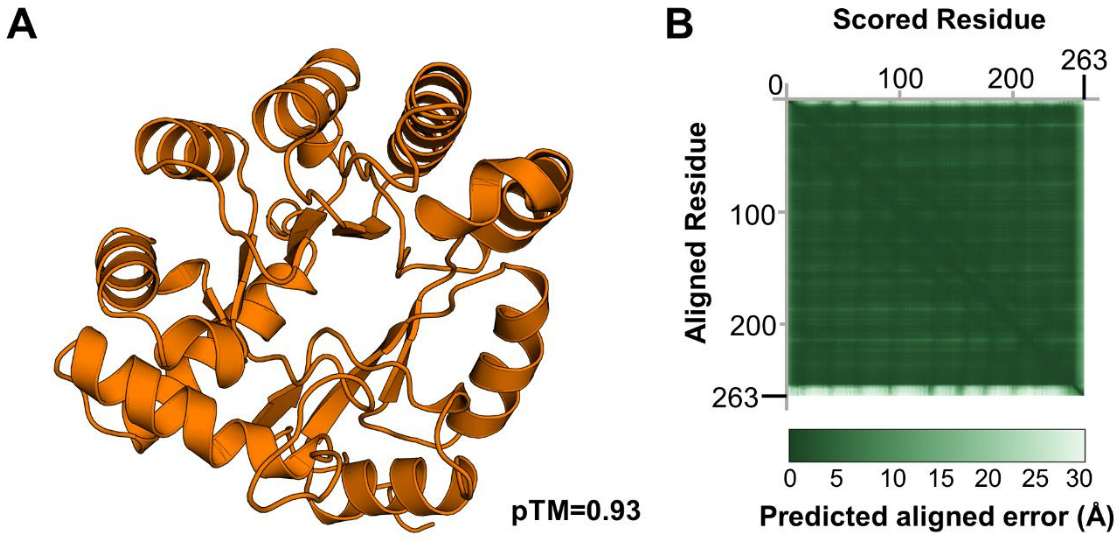


**Figure S9. AlphaFold3 model of QatD**. (A) Predicted structure of QatD generated by AlphaFold 3. The predicted template modelling (pTM) score is provided. (B) Predicted aligned error plot of the QatD model.

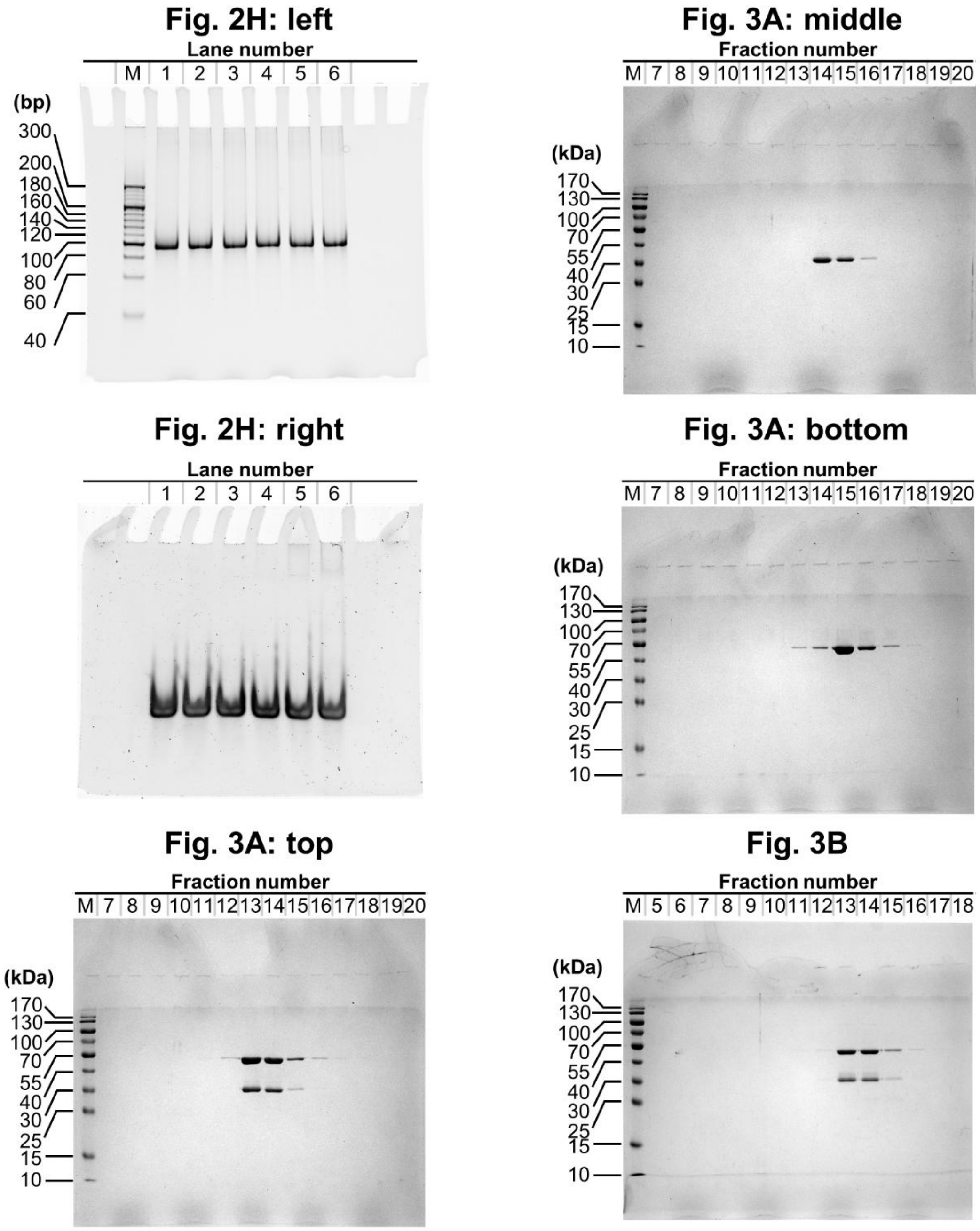


**Figure S10. Uncropped gel images.**

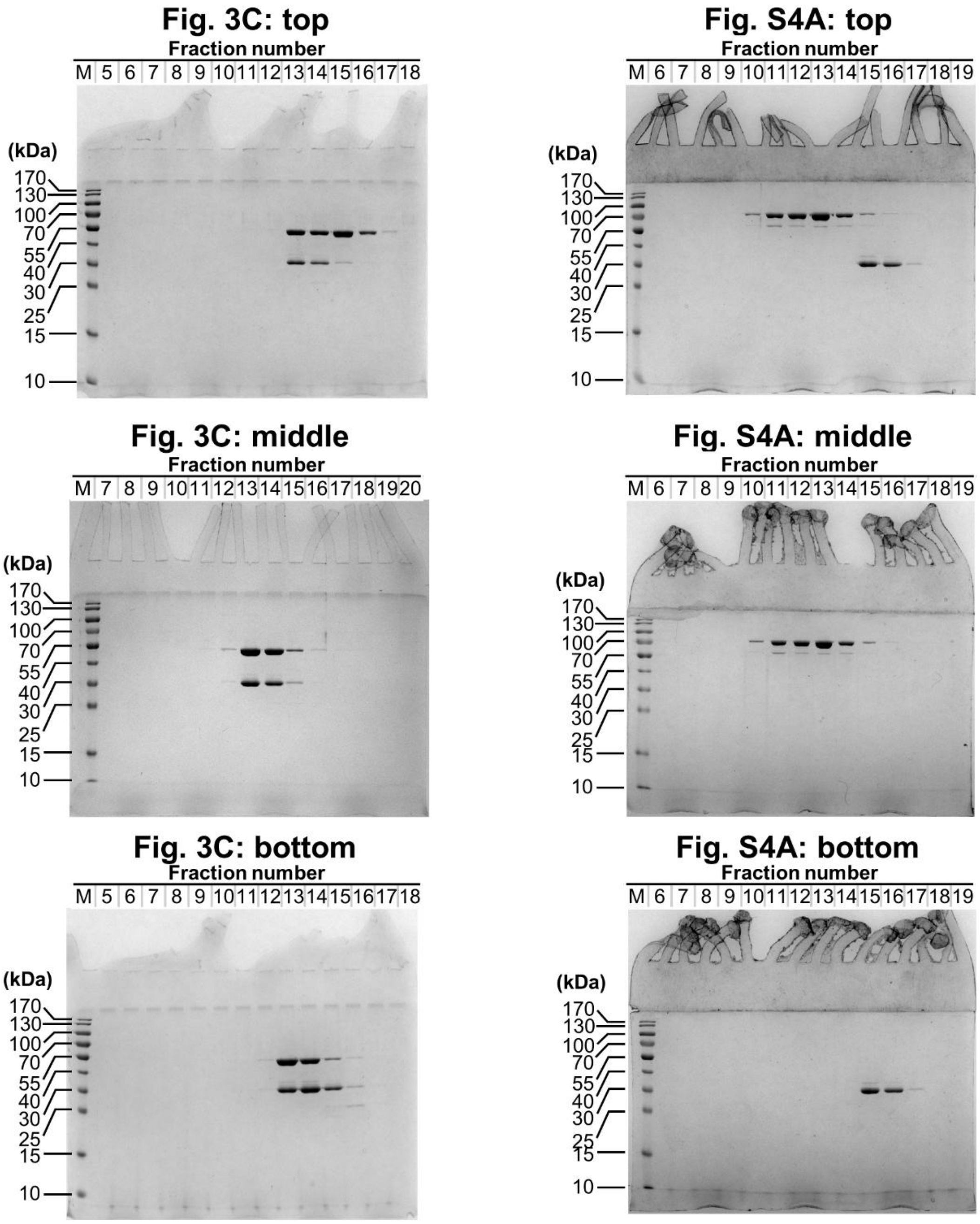


**Figure S10. Uncropped gel images (continued).**

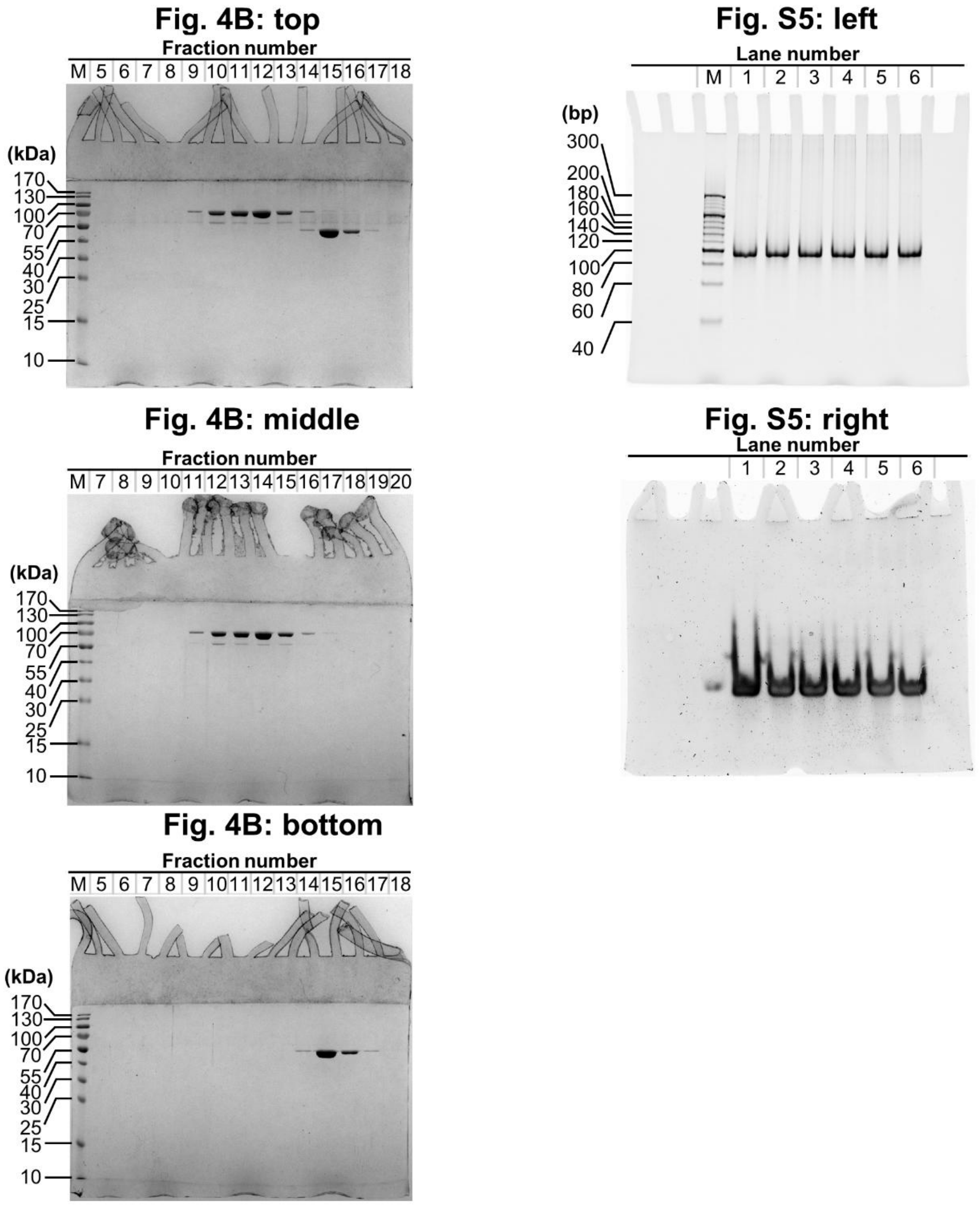


**Figure S10. Uncropped gel images (continued).**
